# Evolution of chronic lymphocytic leukemia after allogeneic stem cell transplant

**DOI:** 10.1101/2025.01.22.634212

**Authors:** Haven R. Garber, Danwei Wang, Celine Kerros, Hannah C. Beird, Xizeng Mao, Nina G. Howard, Jianhua Zhang, Jason Roszik, John P. Miller, Paul Leonard, Yu Cao, Li Zhao, Xingzhi Song, Sahil Seth, Pei Lin, Huandong Sun, Lisa S. St John, Sijie Lu, William Wierda, Issa F. Khouri, Karen Clise-Dwyer, Jin S. Im, Gheath Alatrash, P. Andrew Futreal, Shoudan Liang, Priya Koppikar, Shengqing (Stan) Gu, Jeffrey J. Molldrem

## Abstract

Allogeneic stem cell transplant (alloSCT) for patients with relapsed/refractory chronic lymphocytic leukemia (CLL) can result in cure in some patients. Analogous to chemotherapy and targeted therapy, we hypothesized that allogeneic cellular immunotherapies, including alloSCT and donor lymphocyte infusion (DLI), would impact malignant evolution through the application of selective immunologic pressure with reciprocal changes in the T cell compartment. We tested a cohort of 24 patients treated with HLA-matched alloSCT +/− DLI, two mediators of the graft versus leukemia (GVL) effect. Comparison of pre-alloSCT samples revealed that a key difference between responders (n=13) and non-responders (n=11) is the cellularity of leukemic cells. We further mapped mutational trajectories of tumor cells by whole exome sequencing (WES) of sort-purified CLL in 11 post-transplant relapsed patients and found evidence of subclonal leukemic evolution in 8/11 patients after nonmyeloablative human leukocyte antigen (HLA)-matched alloSCT. Different patterns of CLL evolution were observed, and these changes included putative CLL drivers in every case. To investigate the presence of immune-related variants in patients, we collected 19 T cell co-culture CRISPR datasets and identified the top positive and negative regulators of cancer cell’s response to T-cell-dependent killing. We found that most mutations linked to T-cell killing emerged after allo-SCT treatment, suggesting that selective pressures from the GVL effect may drive the evolution of these mutations. Together, these data identify cellular homogeneity as a key biomarker for susceptibility to GVL-driven immunity in CLL and illustrate how the leukemic cells further evolve to evade immunosurveillance.

**SIGNIFICANCE:** The impact of allogeneic stem cell transplant (alloSCT) on subclonal leukemia evolution remains poorly understood. By performing whole exome sequencing (WES) on pre- and post-alloSCT patient samples, we reveal different patterns of CLL evolution and potential immune-related driver genes influencing CLL relapse and refractory disease post-alloSCT.

## INTRODUCTION

Chronic Lymphocytic Leukemia (CLL) is a malignancy of mature B lymphocytes that demonstrates significant clinical heterogeneity. A majority of patients experience an indolent disease course, permitting them to live for years without treatment. However, about a third of patients experience a rapid progression of disease requiring immediate intervention, with many experiencing chemo-refractory disease (1). Understanding tumor characteristics that contribute to this clinical variation is essential to the development of personalized treatment and improving patient outcomes.

Intratumoral heterogeneity is one such feature that has been repeatedly implicated in limiting the durability of cancer therapies across solid tumor and pan-cancer studies (2). Treatment-driven selection of subclones within a tumor population can cause a patient’s malignancy to evolve, often leading to more aggressive disease (3). To improve the durability of cancer therapies and to rationally design combinations, it is important to understand how various treatments impact malignant evolution.

In CLL, the standard of care has been transformed by the use of targeted therapies such as kinase inhibitors (4). However, allogeneic stem cell transplantation (alloSCT) remains the only cure to the disease (5). This is particularly important in patients with aggressive phenotypes, who are often refractory to available first-line treatment regimens. The efficacy of alloSCT relies on donor T cells to eliminate leukemia through the recognition of tumor antigens – the Graft versus Leukemia (GVL) effect. A related cellular therapy is donor lymphocyte infusion (DLI), which is used to treat post-alloSCT relapse. The evolution of leukemia after chemotherapy and targeted therapies has been investigated (6-11), however, the impact of alloSCT on subclonal evolution is largely unknown. Furthermore, evolution in myeloid malignancies has been more often studied as stem cell transplant is more readily performed, but the impact of transplant on CLL is less understood (12).

Landau et al. reported leukemic evolution in a majority of patients with CLL (57 of 59) after chemotherapy and noted therapeutic selection of subclones with TP53 mutations and del(17p) (10). CLL relapse in patients treated with ibrutinib was accompanied by the outgrowth of clones that expressed the canonical resistance mutation BTK (C481S), as well as mutations in its target protein encoded by PLCG2, and del(8p) (6). We sought to investigate the evolutionary pressure imposed by alloSCT and DLI and hypothesized that allogeneic T cells mediating GVL could shape the evolution of CLL. Here, we test this prediction utilizing whole exome sequencing (WES) of sort purified leukemia cells from a cohort of 24 patients treated with alloSCT and DLI, of which 13 experienced complete responses (denoted as complete responders; CR). Our cohort also included 11 patients who relapsed within 2 years after alloSCT (denoted as non-responders; NR), and in these patients, we sampled longitudinal time points to examine leukemic evolution.

By comparing the leukemic landscape in complete-responders and relapsed patients, as well as the clonal and subclonal evolution following transplant, we are able to elucidate important tumor intrinsic characteristics that can portend success in stem cell transplant. We present factors which may contribute to GVL resistance, which has important implications for therapeutic algorithms, and is essential to understand in relapsed and refractory disease.

## RESULTS

### Patient Characteristics

We analyzed available blood and bone marrow leukemia samples from 24 patients who received non myeloablative human leukocyte antigen (HLA)-matched alloSCT for CLL at MDACC between 1999 and 2009. Our patient cohort was relatively young at the time of CLL diagnosis (median age 53 years) because younger patients tend to demonstrate more aggressive disease (13) and are more fit for transplant (**Table S1**). All patients were heavily pre-treated and demonstrated resistance to frontline chemotherapy with fludarabine, cyclophosphamide plus rituximab (FCR) prior to alloSCT, and all but 1 of the 24 patients received nonmyeloablative (NMA) or reduced-intensity conditioning (RIC), which relies on GVL for curative effect (**Table S2**). Unmutated immunoglobulin heavy-chain variable region gene (*IGHV*) is among the strongest adverse risk factors in CLL (14). Notably, all 21 patients tested had unmutated *IGHV* (3 patients had insufficient sample). CLL cells (CD19+CD5+) were purified from cryopreserved BM and PB samples prior to DNA extraction. We performed WES to an average depth of 120X to detect exonic somatic single nucleotide variants (sSNVs) in the leukemia. We detected a mean of 41.2 (SD ± 15) sSNVs (exonic silent, non-silent, and indels) per case (**Fig. 1A**).

**Fig. 1.**
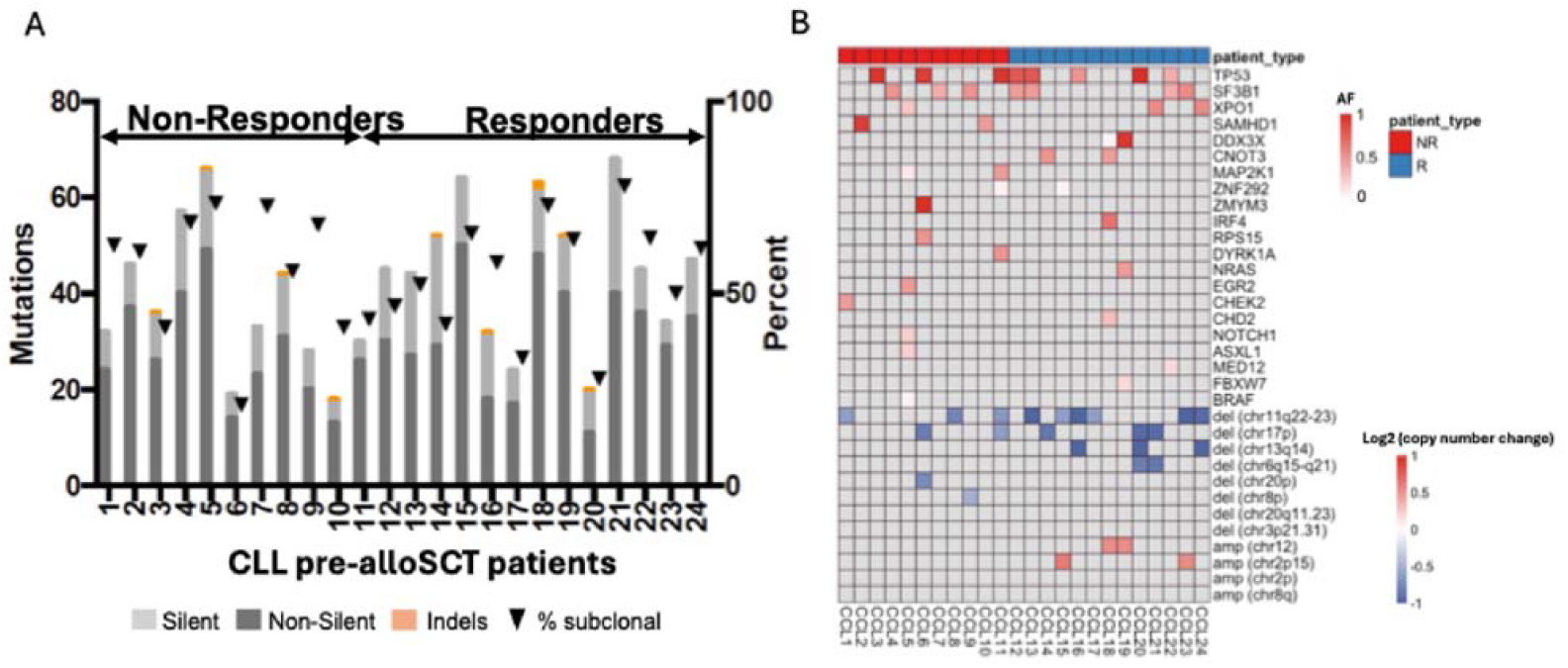
Patient Characteristics. **A**, Twenty-four patients with CLL were enlisted in this study. Total numbers of somatic exonic silent and non-silent SNVs and indels are listed for each patient’s pre-alloSCT leukemia (allelic fraction (AF) > 0.05). The percentage of mutations that are subclonal is also indicated (triangle). **B**, The landscape of somatic variants in genes and chromosome regions recognized to be CLL drivers is shown for the CLL alloSCT cohort. Variants are shaded according to their corresponding variant AF or exome-derived copy number log2 ratio.

The patients in our cohort had leukemia enriched with variants in recognized CLL drivers (10, 15) (**Fig. 1B**). The mutation for TP53 and chromosome deletion for chr11q are found in most of patients. A median of 3 drivers genes was observed per case, and most of patients harbored 3 or more drivers prior to alloSCT.

### Differences between Responder and Non-responder patients

To compare the differences between the non-responder and responder patients, we first used WES data to infer tumor heterogeneity based on the number of clonal and subclonal mutations. There are no significant differences in the number of subclonal mutations or clonal mutations between pre_AlloSCT non-responder patients and responder patients (**Fig. 2A and 2B**). Considering the potential impact for chromosomal aberrations, ‘cellularity’ was also performed here. Cellularity, as defined in Sequenza, is the fraction of tumor cells in a patient sample. Sequenza derives the cellularity parameter from WES from 2 input genomes: normal cells (for germline analysis) and the tumor sample, which is assumed to be a mixture of normal and tumor cells. The program determines copy-number variation and allele fraction (AF) at SNP sites across the genome and compares tumor and normal samples region by region to find the most likely cellular fraction (with values ranging between zero to one, a value of one indicates a purely clonal tumor sample). Somatic mutated alleles are not used in the calculation. Since we used pre-transplant sorted leukemia cells for these analyses, we postulated that when cellularity is close to one, the tumor sample was pure according to copy number and SNP heterogeneity. Accordingly, when the cellularity is less than one, it meant the tumor sample was comprised of at least two structural clones. Our results showed that responder cohort showed a singular pattern of tumor cellularity for alloSCT treatment (**Fig. 2C)**, which means responder cohort tended to be more structurally clonal and has more homogenous copy number gain or loss. To understand whether the higher cellularity observed in CR patients was limited to our alloSCT cohort, we carried out similar analyses on CLL samples collected before and after relapse following chemotherapy from previously published data sets (10). We examined tumor cellularity of pretreatment samples from 10 randomly selected relapsed patients (out of 59 evaluable patients) and compared them to pretreatment samples from randomly selected 15 patients that went into remission following FCR therapy (from a pool of 219 evaluable patients). Sequenza analysis did not find any statistically significant differences between these two cohorts (**Fig. 2D)**, which means the level for cellularity might be the potential predictor for the allo-SCT treatment outcome but not chemotherapy. We also predicted potential neoantigens for each patient in these two groups and found no significant differences in the number of predicted total or strong-binding neoantigens (**Fig. 2E and 2F)**.

**Fig. 2.**
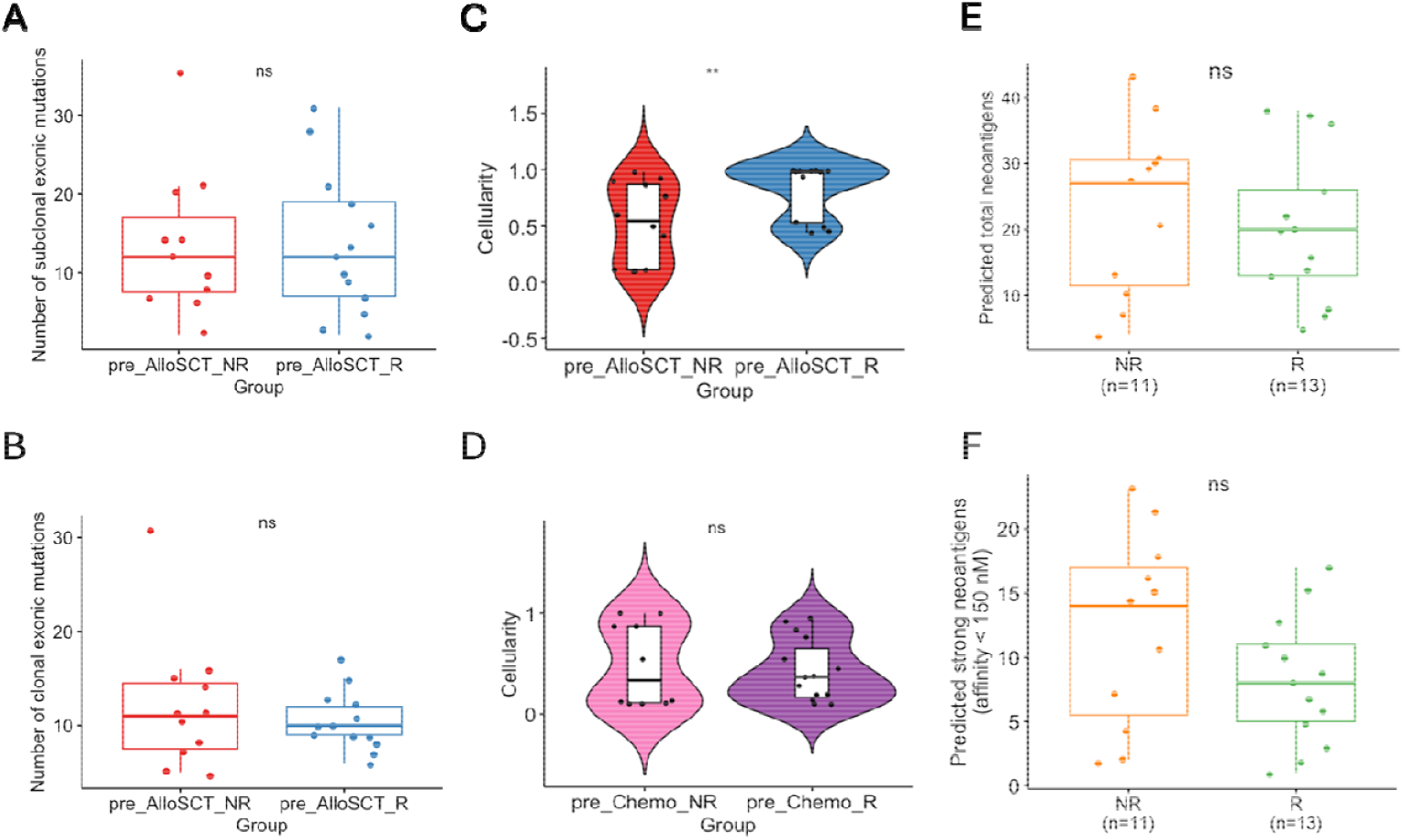
Differences between Responder and Non-responder patients. **A**, Comparison of clonal exonic mutations (p=0.84, Wilcoxon rank sum test) between the transplant early relapse and durable CR groups indicates no significant differences between the two groups of patients. **B**, Comparison of subclonal exonic mutations (inset; p=0.82, Wilcoxon rank sum test) between the transplant early relapse and durable CR groups indicates no significant differences between the two groups of patients. **C**, Cellularity of pre-transplant CLL samples was determined by Sequenza v3.0. WES data from sort-purified CLL cells were analyzed, showing lower clonality in NR patients compared to high clonality in CR patients (p=0.0078, Wilcoxon rank sum test). **D**, In pre-treatment samples, cellularity differences between chemo responders (n=15) and non-responders (n=10) were not significant (p=0.87). **E**, Comparison of the total (p=0.58, Wilcoxon rank sum test) neoantigen burden between CLL transplant early relapse and durable responder groups indicates no significant differences between the two groups. **F**, Comparison of strong binding (inset; p=0.19, Wilcoxon rank sum test) neoantigen burden between CLL transplant early relapse and durable responder groups indicates no significant differences between the two groups.

### Different patterns of evolution in the Non-responder patients

To examine leukemic evolution after alloSCT, we compared the AF of somatic mutations detected in longitudinal CLL patient samples. Clear patterns of evolution emerged, and in 2 patients (CLL 5 and CLL 8), evolution coincided with DLI administration for relapsed disease. The clinical course and WES sampling windows are shown for CLL 5 and CLL 8 (**Fig. 3A and 3B)**. Sort purified CLL was sequenced from 5 time points from both sets of samples (**Figs. 3C and 3D**). Clinical course and WES windows for remaining 22 CLL patients are shown in (**Fig. S1)**. Branched leukemic evolution was seen with a branch point between time points 03/2005 and 04/2006, when CLL 5 received 3 DLIs (**Fig. 3C)**. Outgrowth of a leukemic subclone containing non-silent mutations in *EGR2, NOTCH1, XPO1*, and a new mutation in *ASXL1* (p.M1345V) was seen post-DLI concurrent with the elimination of a subclone containing *NOTCH1* nonsense mutation (p.S2492X) among several other variants. CLL 8 experienced a similar relapsing and remitting clinical course (**Fig. 3B**), and branched leukemic subclonal evolution again coincided with DLI treatment (**Fig. 3D**). The subclone ASXL1(p.W1037X) and TP53(p.G244A) were expanded post-DLI. We performed this analysis in all 11 patients with available longitudinal post-alloSCT samples and observed branched CLL evolution in 5 patients (**Fig. S2A**), linear evolution in 3 patients (**Fig. S2B**) and no evolution in 3 patients (**Fig. S2C**). Notably, studies of CLL evolution after chemotherapy have also demonstrated mixed patterns of branched and linear evolution (9, 10). The data support our hypothesis that allogeneic T cells can shape leukemic subclonal architecture after transplant.

**Fig. 3.**
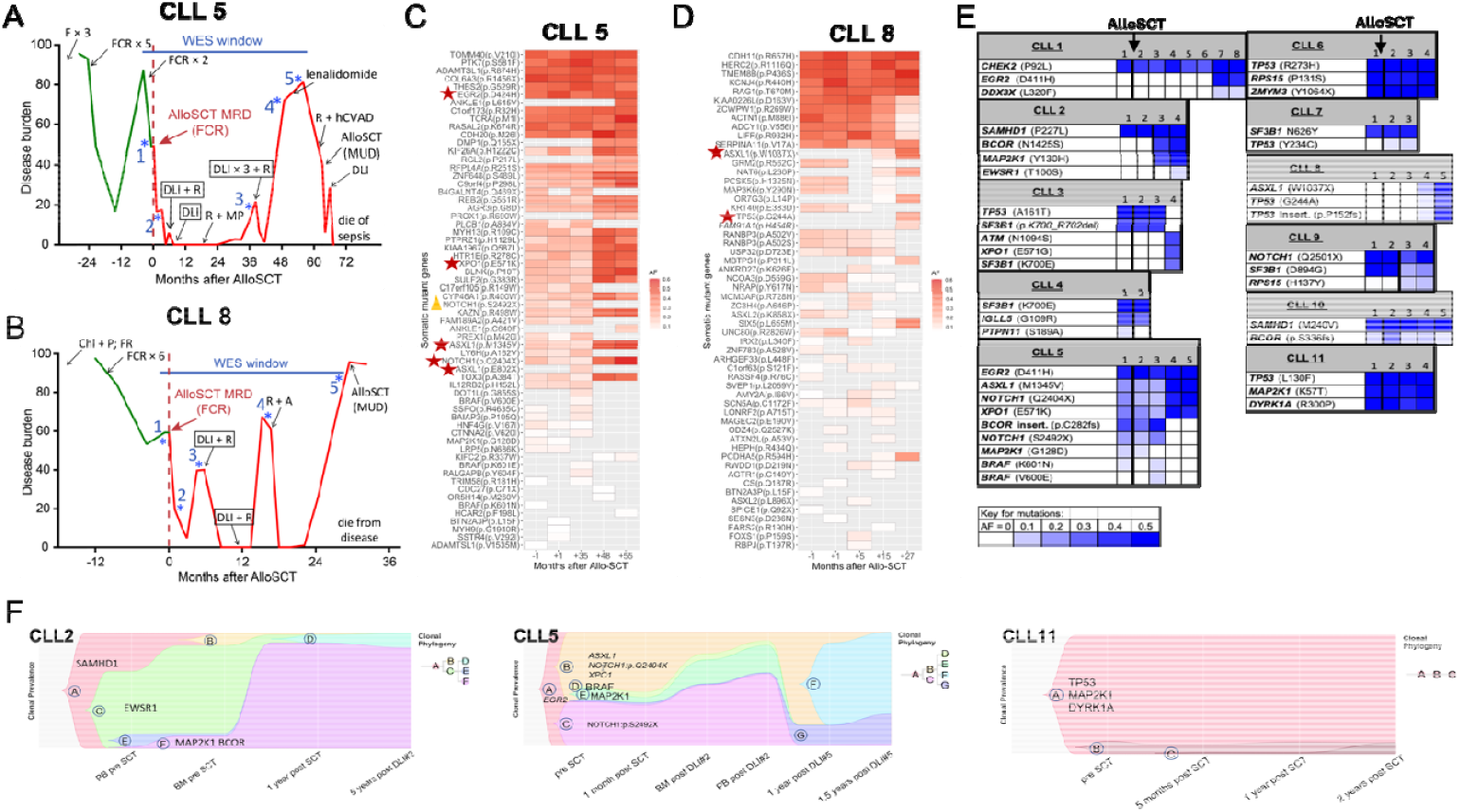
Different patterns of evolution in the Non-responder patients. **A**, Clinical course for CLL 5. Green lines indicate disease burden prior to an alloSCT, red lines indicate disease burden following transplant. **B**. Clinical course for CLL 8. Green lines indicate disease burden prior to an alloSCT, red lines indicate disease burden following transplant. **C**, Heatmap indicating shifts in reported CLL driver lesions over time for CLL5 patient, shaded by AF (red). **D**, Heatmap indicating shifts in reported CLL driver lesions over time for CLL8 patient, shaded b AF (red). **E**, The 11 NR patients with longitudinal samples are shown with driver gene mutations, shaded by AF (blue). Each column represents a longitudinal time point in a patient’s course and the bar above indicates the number of years between the first and last WES time point. Solid bars in bold indicate interval alloSCT (indicated by arrows). **F**, CALDER analysis of CLL 2, CLL 5 and CLL 11 tumor evolution post alloSCT. WES data from sort purified CLL was used for the analyses. Pipeline included Pyclone analysis followed by Mixed integer Linear programming (using Gurobi engine) and visualization by Timescape. The X axis for each sample indicates time points when WES analysis was performed. Y axis indicates clonal prevalence. Clonal phylogenetic trees are indicated to the right of each figure. (The meaning of the abbreviations in **A & B** is shown in **Table S2**.)

We anticipated there would be immunoediting of the leukemic subclones by allogeneic T cells; however, given that most CLL subclonal selection had likely already resulted from prior treatments (10), it was unknown whether relapsed CLL post-alloSCT and DLI would differ substantially from the pre-alloSCT leukemia, particularly with regard to established driver lesions. To answer this question, we looked for changes in CLL drivers over time in the 11 patients with sequential samples available post-transplant. We observed marked evolution of variants in established CLL driver genes and chromosome regions after alloSCT (**Fig. 3E**). For example, CLL 2 demonstrated linear evolution of CLL post-alloSCT, and the relapsed leukemia contained mutations in the drivers *BCOR* and *MAP2K1* (16) as well as a mutation in *EWSR1* (p.T100S). CLL 6 also demonstrated linear evolution post-alloSCT and acquired an 18p deletion late after treatment. After alloSCT, 44% of CLL drivers remained unchanged in their AFs; 36% expanded, and 20% contracted. Six of 11 patients saw the emergence of at least one previously undetectable driver post-transplant, including mutations in *EGR2, XPO1, SF3B1*, and *TP53* as well as mutations in *DDX3X* (p.L320F) and *ATM* (p.N1094S). Our data suggest that CLL evolution continues to be dynamic post alloSCT. For a formal analysis of ancestral relationship between CLL clones before and several time points after transplant, we applied the program, CALDER (Cancer Analysis of Longitudinal Data through Evolutionary Reconstruction) (17). CALDER has been used to infer tumor phylogeny in longitudinal CLL samples by converting somatic SNV clusters into clones (17). We adapted a similar strategy by using Pyclone (18) for statistical inference of clonal populations followed by CALDER analysis. As predicted, our results suggest that CLL cells are subject to immune selective evolution post alloSCT (CLL 2 and CLL 5), consistant with immunoediting. CLL 11 is shown as an example where disease did not evolve despite intervening therapy and sub clinical response (**Fig. 3F**).

### Mutations in genes associated with response to T-cell killing in the non-responder and responder patients

To investigate the presence of immune-related variants in patients, we collected 19 T cell co-culture CRISPR datasets from 9 published studies and identified the top positive and negative regulators of cancer cell response to T-cell-dependent killing. Positive regulators are defined as genes that enhance the sensitivity of cancer cells to T-cell killing, while negative regulators are genes that inhibit this sensitivity. We investigated the mutation patterns of these positive and negative regulators in pre-AlloSCT responder and non-responder patients. Additionally, we examined the evolutionary dynamics of these regulators at different time points in non-responder patients.

In our pre-AlloSCT patient cohort, only a small fraction of patients harbored mutations in genes linked to response to T-cell killing (**Fig. 4A and 4B**). Notably, 10 out of 11 non-responder patients acquired mutations in these genes during post-transplant evolution. (**Fig. 4C and 4D**), suggesting that selective pressures from the transplant environment may drive the evolution of these mutations.

**Fig. 4.**
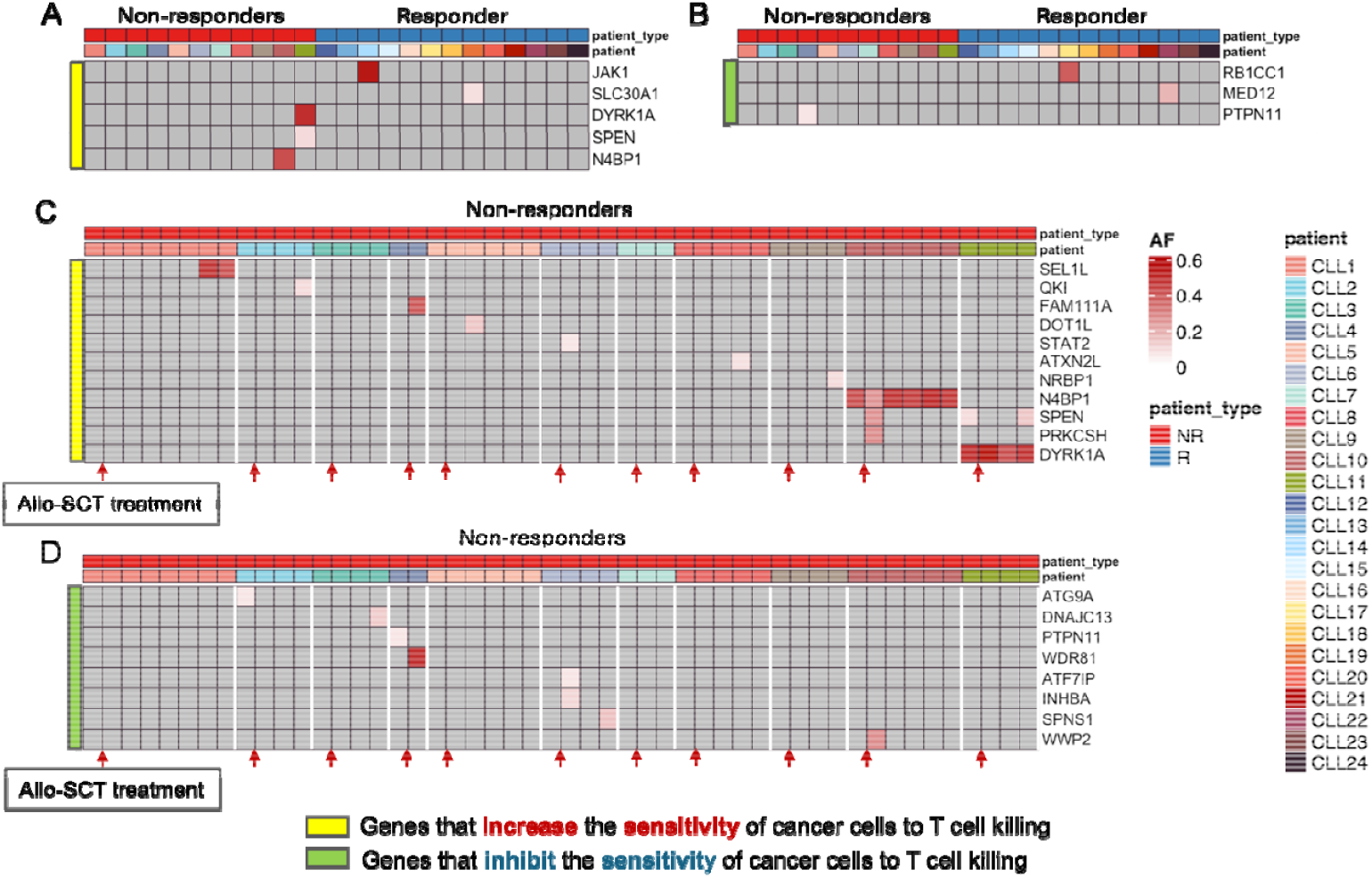
T-cell killing-associated mutations in the non-responder and responder patients. **A**, Genes that increase the sensitivity of cancer cells to T cell killing in pre-AlloSCT non-responder and responder patients, shaded by AF (AF > 0.05). **B**, Genes that inhibit the sensitivity of cancer cells to T cell killing in pre-AlloSCT non-responder and responder patients, shaded by AF (AF) > 0.05). **C**, Mutation pattern of these genes that increase the sensitivity of cancer cells to T cell killing at different time point in NR patients, red row is the time point for patients receiving AlloSCT treatment. The upper color bar corresponds to the different non-responder patients. **D**, Mutation pattern of these genes that inhibit the sensitivity of cancer cells to T cell killing at different time point in NR patients, red row is the time point for patients receiving AlloSCT treatment. The upper color bar corresponds to the different non-responder patients.

## DISCUSSION

Here, we demonstrate that tumor cellularity was the singular predictor of outcome to stem cell transplant. The SCT responder cohort was more structurally clonal than non-responders, and this distinction was not seen following chemotherapy. This suggests cellularity of leukemic cells prior to transplant may be a unique prognostic marker for allogeneic stem cell transplant response, and that therapeutic strategies should account for this distinction. Indeed, very few commonly mutated genes were identified in the non-responder patients, reflecting the high degree of cancer heterogeneity among this cohort. While studies of clonality in CLL patients undergoing SCT have previously been limited, tumor heterogeneity has been repeatedly implicated as an important predictor of immune response and patient outcome in pan-cancer studies. Our data provide important evidence that tumor heterogeneity is a key modulator of graft-versus-leukemia resistance in CLL.

We also demonstrate that CLL evolution continues to be dynamic post-transplant, and that subclone architecture is shaped by transplant. Among subjects post allo-SCT relapse, marked evolution of variants in established driver genes was shown, with the emergence of previously undetectable driver mutations in six of eleven patients. In addition, distinct subclonal evolutionary patterns were observed in this transplant relapsed cohort.

Finally, we demonstrate that most mutations in genes associated with T cell killing arose after SCT among patients who relapsed following transplant. As the efficacy of alloSCT relies on donor T cells’ elimination of leukemia through the recognition of tumor antigens, the GVL effect, understanding how CLL responds to the T cell impact is essential to understand how GVL resistance may evolve. Our data suggest that GVL effect of alloSCT shapes the evolution of CLL post-transplant.

We provide a tumor cell-intrinsic characteristic; structural clonality, that can potentially predict response to alloSCT. This data supports the model proposed by George et al, which predicted lower clonal heterogeneity would favor susceptibility to immune responses (19). While our tumor clonality data are intriguing, we note that our small patient sample size may not be reflective of tumor clonality association with transplant outcomes in a larger patient cohort, and that any potential association between tumor clonality and tumor remission post alloSCT needs to be evaluated in a larger sample size.It would also be interesting to understand if these findings are translatable across malignancies, especially in AML where the disease course is accelerated.

In conclusion, we present a tumor intrinsic characteristic, allelic cellularity, that can portend treatment success using stem cell transplant in CLL. We demonstrate that there is ongoing CLL subclonal evolution after HLA-matched alloSCT, including known CLL drivers, among patients who relapse following transplant. We also observe prevalent mutations in genes associated with response to T-cell killing in non-responders following alloSCT. These findings bring new insight to the underlying mechanisms of alloSCT resistance in CLL and the prediction of treatment response.

## METHODS

### Study Design

The main objective of our retrospective analysis was to evaluate the interaction of leukemia with the post-transplant adaptive immune cells. We chose twenty-four patients with a confirmed diagnosis of CLL based on NCCN guidelines and flow cytometry analyses for our analyses. All patients received an allogeneic stem cell transplant at MDACC between 1999 to 2009. Patient and healthy donor samples were obtained after appropriate informed consent through institutional review board approved protocols at The University of Texas MD Anderson Cancer Center (MDACC). Those patients in the transplant responder subset experienced a CR after alloSCT as defined in the guidelines set by the International Workshop on CLL (20). Eleven patients relapsed after transplant (denoted as non responders, NR) and in these patients, we performed WES to analyze leukemic evolution. Neoantigen prediction was performed to correlate tumor subclonal kinetics with antigen specific Graft versus Leukemia (GVL) T cells. The study was not blinded, and biological replicates were not performed, given limited sample availability.

### Patient samples

Patient peripheral blood (PB) and bone marrow (BM) mononuclear cells were separated using Histopaque 1077 (Sigma-Aldrich) prior to initial cryopreservation (FBS with 10% DMSO) and were stored in vapor phase of liquid nitrogen until the time of analysis. All patients received a matched alloSCT for CLL at MDACC. Overall, we sequenced 76 leukemia samples (68 sort purified PB/BM, 5 unsorted BM, and 3 formalin-fixed paraffin embedded (FFPE) BM) and 24-matched germline samples (primarily sort purified T cells) from 24 patients along with 6 matched alloSCT donor samples. Serial samples were sequenced in 20 of 24 patients to validate the detected variants.

HLA status and clinical parameters, including molecular studies, cytogenetics, prior treatments, and outcomes were obtained from the patients’ electronic medical record. All patients in the CLL evolution subset (n = 11, NR) experienced disease relapse post-alloSCT, which was confirmed by pathology and flow cytometry. For 20/24 patients, HLA testing was conducted at the MDACC HLA typing laboratory. For 4 patients with only serologic HLA typing available, HLA typing was inferred from exome data using the winners output from Polysolver (21). BM and PB samples were collected prior to transplant and at time of disease progression/response after alloSCT.

### Cell purification

PB mononuclear cells and BM aspirate cells were thawed and stained with the following antibodies prior to electrostatic droplet-based cell sorting: anti-CD19 FITC (clone SJ2SC1), anti-CD5 PE (clone UCHT2), anti-Ig l light chain Pacific Blue (clone MHL-38), anti-Ig k light chain Pacific Blue (clone MHL-49), anti-CD3 PE/Cy7 (clone SK7), anti-CD8 PE/Cy7 (clone SK1) (all from BioLegend) and Sytox Red live/dead stain (ThermoFisher). CD19+CD5+ CLL cells and CD3+ T cells were sorted on the FACSAria Fusion (BD Biosciences) using a 70 μM nozzle at 70 psi with a purity mask (Y32-P32-Ph0) after excluding debris, doublets, and dead cells. Cells were thawed and stained in phosphate-buffered saline containing 2% FBS. Sorted cells were collected in RPMI before DNA isolation.

### Whole exome sequencing

Genomic DNA was extracted from CLL cells and pre-alloSCT T cells (for germline DNA) using the QIAamp DNA Blood Mini Kit (Qiagen). Three FFPE BM aspirate samples were used for analysis. DNA was extracted from the FFPE aspirate blocks using the QIAamp DNA FFPE Tissue kit (Qiagen). Tumor and germline DNA concentration and quality were measured using fluorometric quantification (Qubit, ThermoFisher, and Fragment Analyzer, Advanced Analytical).

Libraries were constructed from genomic DNA using the KAPA Library Preparation Kit (Roche). Exome capture was performed using the NimbleGen SeqCap EZ Exome Enrichment Kit v3.0 (Roche). Multiplex sequencing of samples was conducted on the Illumina HiSeq 2000 using 76 base pair paired-end reads at the MDACC Sequencing and Microarray facility. The mean target coverage was 120X per tumor sample (range: 39 – 389; SD +/−45) and 112X per germline sample (range: 50 – 162; SD +/−31).

Paired-end sequencing reads in FASTQ format were generated from BCL raw data using Illumina CASAA software and aligned to the human reference genome (UCSC Genome Browser, hg19) using Burrows-Wheeler Aligner on default settings with the following exceptions: seed length of 40, maximum edit distance of 3, and maximum edit distance in the seed of 2 (22). Aligned reads were then processed using the GATK Best Practices of duplicate removal, indel realignment, and base recalibration. Sequencing was targeted to an overall coverage of 120X for target samples and 100X for matched germline samples.

### Subclonal analysis

The Sequenza package v3.0 (23) was used on the paired tumor-normal BAM (Binary Alignment Map) files to estimate the global parameters of cellularity and ploidy as well as allele-specific copy number alteration profiles. The reads with low-quality mapping were excluded with the parameter *-q30*. Randomly selected samples from previously published FCR refractory/remission CLL WES data sets (10) were downloaded from dbGaP following institutional approval and converted to BAM format using the SRA Toolkit from NCBI.

### Neoantigen prediction

To predict potential neoantigens for each patient, a peptide list was generated from each missense mutation that was identified from the exome sequencing data. Peptides included all possible 9- and 10-mer peptides containing the alternate amino acid that resulted from the missense mutations (24). The binding affinities for each wild type and mutant peptide to the patient-specific HLA molecules (HLA-A and HLA-B) were then tabulated using NetMHCpan (v.2.8). Peptide-HLA complexes with IC50 values < 150 nM were considered strong binding neoepitopes and those with IC50 between150 nM and 500 nM were considered intermediate binding neoepitopes.

### Crispr screen data analysis

To investigate potential immune-related mutations, we collected 33 T cell co-culture CRISPR datasets from 9 published studies. The datasets were analyzed using MAGeCK (25). The top 100 positive and negative regulators of cancer cell response to T-cell-dependent killing were identified according to their respective ranking as positive or negative regulators for subsequent analysis.

### Statistical analysis

Statistical analysis was performed using GraphPad Prism software and R Statistical Language. The Wilcoxon rank sum test was used to compare differences in mutation and copy number data between groups. Two-tailed *P* values were calculated and *P* values of less than 0.05 were considered to be statistically significant. Wilcoxon rank sum test was used to compare pre-transplant alloSCT CLL samples from our cohort as well as the chemo relapsed/responder cohorts analyzed using Sequenza v3.0. For the pre/post FCR samples from the Landau *et al*. data sets (10), paired Wilcoxon signed rank test was used.

## Acknowledgments

The authors are grateful to the patients for their participation. The authors also thank Lauren Hodgson and Rahul Popat for their help with organizing the patient samples for analysis, Rachel Sargent for patient cases and material and Annalea Elwell for her help with formatting the manuscript.

## Funding

Principal funding for the study was provided by the MD Anderson Moon Shots Program (to J.J.M.). The MDACC South Campus Flow Cytometry and Cell Sorting Core and the Sequencing and Microarray Facility are supported by NCI P30CA16672 (to K.C-D).

## Authors’ Contributions

J.J.M., S.S.G. and P.A.F. conceived and designed the study, supervised the experimental work, and edited the manuscript. H.R.G. performed experiments, H.R.G. and D.W. interpreted the data, and wrote the manuscript. D.W and H.C.B. performed bioinformatics analysis and edited the manuscript. C.K., J.P.M., and P.L. performed experiments, interpreted data, and edited the manuscript. X.M. J.Z., J.R., L.Z., X.S., S.S., H.S. and S. Liang performed bioinformatics analysis. Y.C. and S.L. performed experiments and interpreted data. P.Lin., W.W., and I.F.K. contributed patient cases and patient material. L.S.SJ., K.C-D., J.S.I., and G.A. supervised experiments and edited the manuscript. P.K. designed and supervised the experimental work, interpreted the data and co-wrote the manuscript. N.G.H. co-wrote the manuscript.

## Authors’ Disclosures

P.K. holds stock options in Amgen, Inc. The Sequenza analysis (X. M., S. Liang, C.K., G. A., P.K. and J.J.M) and neoantigen sequences (H.R.G., C. K., P. K., X. M., S. Liang, J.J.M.) are part of planned patents. J.P.M. is a current employee at Asylia Therapeutics. All other authors declare no competing interests.

## Data and materials availability

All data are available in the main text or the supplementary materials.

