## Supplementary materials for "Evolution of chronic lymphocytic leukemia after allogeneic stem cell transplant"

### Slide 1
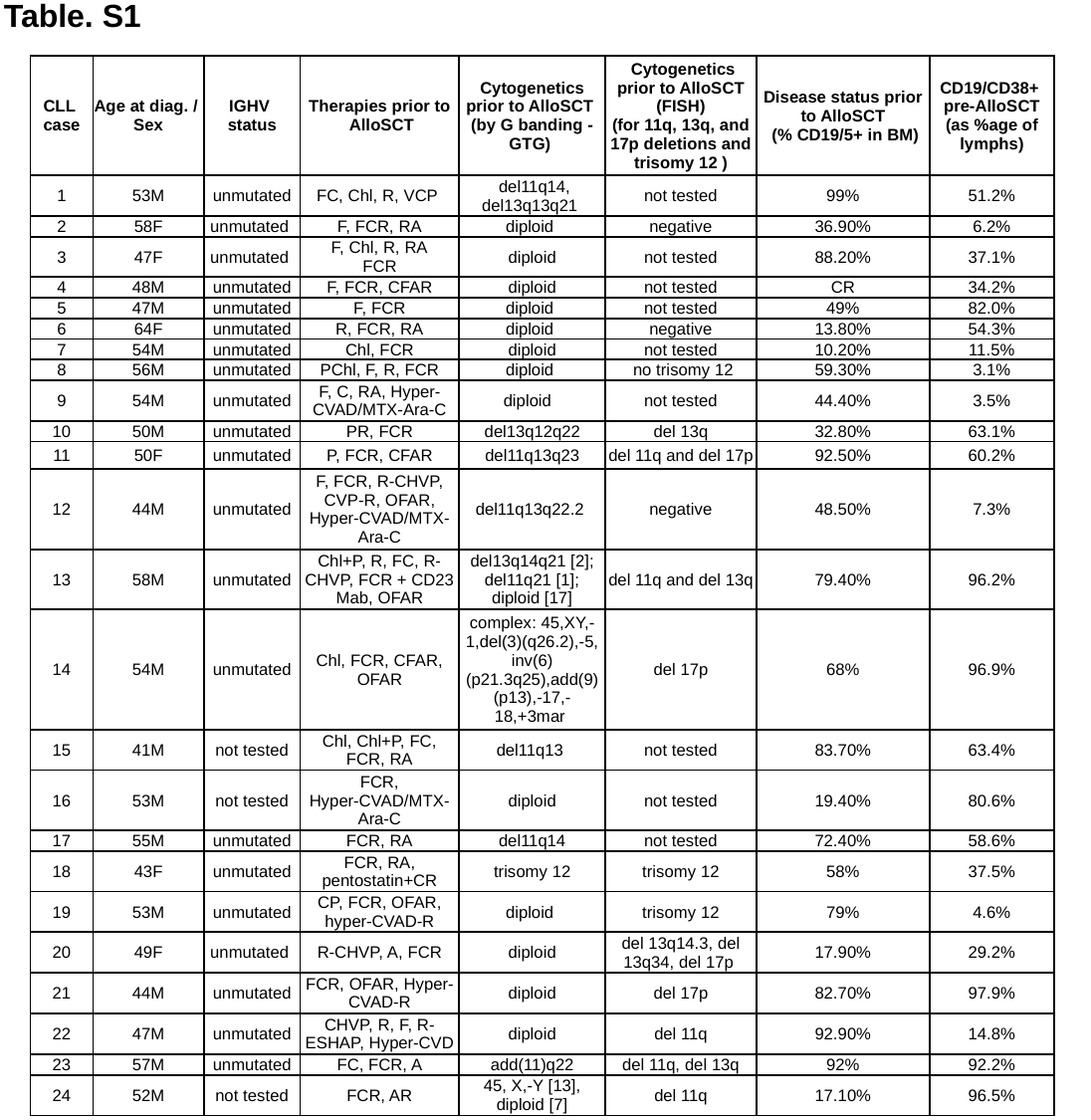

Table. S1
| CLL case | Age at diag. / Sex | IGHV status | Therapies prior to AlloSCT | Cytogenetics prior to AlloSCT (by G banding - GTG) | Cytogenetics prior to AlloSCT (FISH) (for 11q, 13q, and 17p deletions and trisomy 12 ) | Disease status prior to AlloSCT (% CD19/5+ in BM) | CD19/CD38+ pre-AlloSCT (as %age of lymphs) |
| --- | --- | --- | --- | --- | --- | --- | --- |
| 1 | 53M | unmutated | FC, Chl, R, VCP | del11q14, del13q13q21 | not tested | 99% | 51.2% |
| 2 | 58F | unmutated | F, FCR, RA | diploid | negative | 36.90% | 6.2% |
| 3 | 47F | unmutated | F, Chl, R, RA FCR | diploid | not tested | 88.20% | 37.1% |
| 4 | 48M | unmutated | F, FCR, CFAR | diploid | not tested | CR | 34.2% |
| 5 | 47M | unmutated | F, FCR | diploid | not tested | 49% | 82.0% |
| 6 | 64F | unmutated | R, FCR, RA | diploid | negative | 13.80% | 54.3% |
| 7 | 54M | unmutated | Chl, FCR | diploid | not tested | 10.20% | 11.5% |
| 8 | 56M | unmutated | PChl, F, R, FCR | diploid | no trisomy 12 | 59.30% | 3.1% |
| 9 | 54M | unmutated | F, C, RA, Hyper-CVAD/MTX-Ara-C | diploid | not tested | 44.40% | 3.5% |
| 10 | 50M | unmutated | PR, FCR | del13q12q22 | del 13q | 32.80% | 63.1% |
| 11 | 50F | unmutated | P, FCR, CFAR | del11q13q23 | del 11q and del 17p | 92.50% | 60.2% |
| 12 | 44M | unmutated | F, FCR, R-CHVP, CVP-R, OFAR, Hyper-CVAD/MTX-Ara-C | del11q13q22.2 | negative | 48.50% | 7.3% |
| 13 | 58M | unmutated | Chl+P, R, FC, R-CHVP, FCR + CD23 Mab, OFAR | del13q14q21 [2]; del11q21 [1]; diploid [17] | del 11q and del 13q | 79.40% | 96.2% |
| 14 | 54M | unmutated | Chl, FCR, CFAR, OFAR | complex: 45,XY,-1,del(3)(q26.2),-5, inv(6)(p21.3q25),add(9)(p13),-17,-18,+3mar | del 17p | 68% | 96.9% |
| 15 | 41M | not tested | Chl, Chl+P, FC, FCR, RA | del11q13 | not tested | 83.70% | 63.4% |
| 16 | 53M | not tested | FCR, Hyper-CVAD/MTX-Ara-C | diploid | not tested | 19.40% | 80.6% |
| 17 | 55M | unmutated | FCR, RA | del11q14 | not tested | 72.40% | 58.6% |
| 18 | 43F | unmutated | FCR, RA, pentostatin+CR | trisomy 12 | trisomy 12 | 58% | 37.5% |
| 19 | 53M | unmutated | CP, FCR, OFAR, hyper-CVAD-R | diploid | trisomy 12 | 79% | 4.6% |
| 20 | 49F | unmutated | R-CHVP, A, FCR | diploid | del 13q14.3, del 13q34, del 17p | 17.90% | 29.2% |
| 21 | 44M | unmutated | FCR, OFAR, Hyper-CVAD-R | diploid | del 17p | 82.70% | 97.9% |
| 22 | 47M | unmutated | CHVP, R, F, R-ESHAP, Hyper-CVD | diploid | del 11q | 92.90% | 14.8% |
| 23 | 57M | unmutated | FC, FCR, A | add(11)q22 | del 11q, del 13q | 92% | 92.2% |
| 24 | 52M | not tested | FCR, AR | 45, X,-Y [13], diploid [7] | del 11q | 17.10% | 96.5% |

### Slide 2
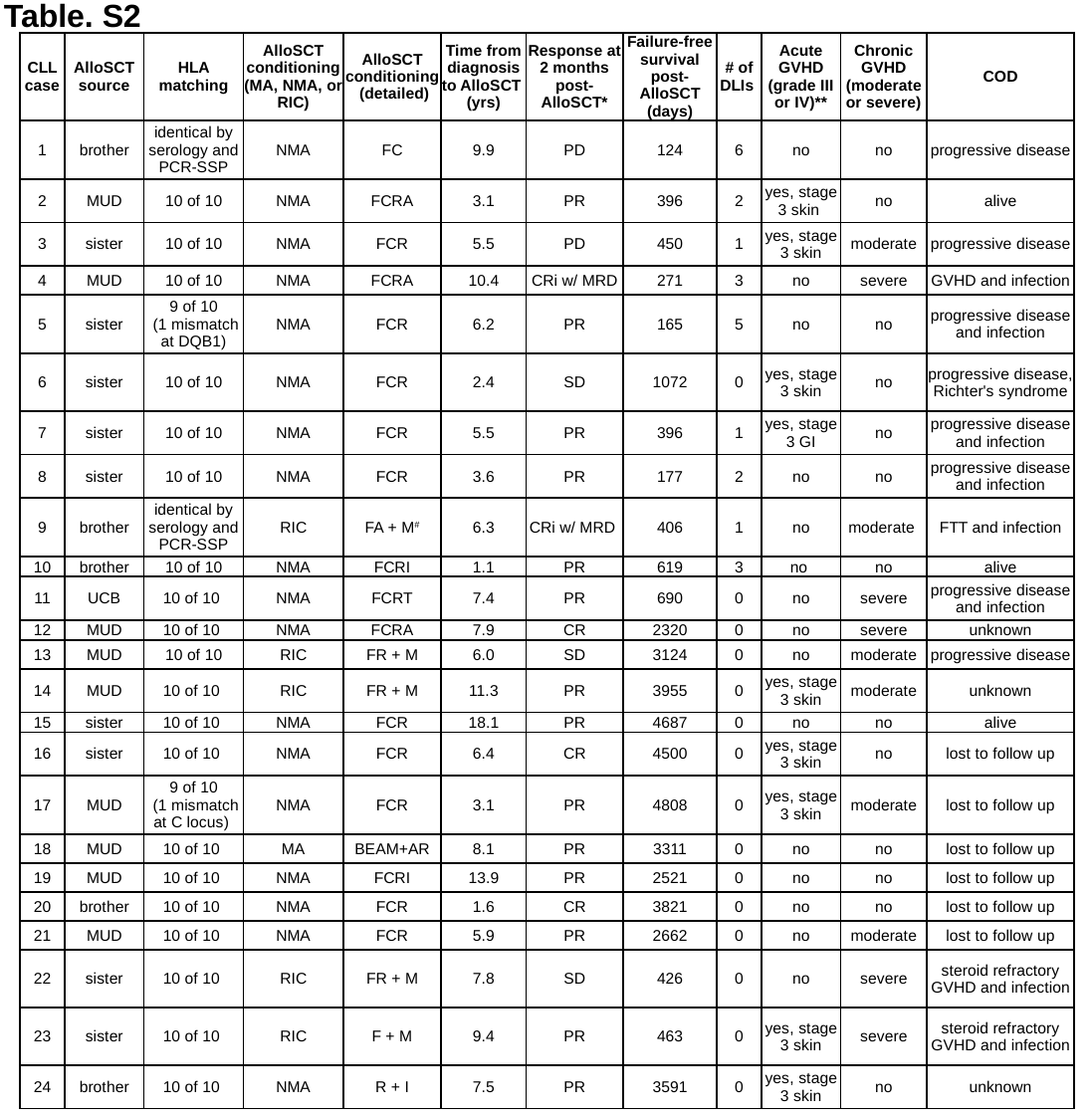

Table. S2
| CLL case | AlloSCT source | HLA matching | AlloSCT conditioning (MA, NMA, or RIC) | AlloSCT conditioning (detailed) | Time from diagnosis to AlloSCT (yrs) | Response at 2 months post-AlloSCT\* | Failure-free survival post-AlloSCT (days) | # of DLIs | Acute GVHD (grade III or IV)\*\* | Chronic GVHD (moderate or severe) | COD |
| --- | --- | --- | --- | --- | --- | --- | --- | --- | --- | --- | --- |
| 1 | brother | identical by serology and PCR-SSP | NMA | FC | 9.9 | PD | 124 | 6 | no | no | progressive disease |
| 2 | MUD | 10 of 10 | NMA | FCRA | 3.1 | PR | 396 | 2 | yes, stage 3 skin | no | alive |
| 3 | sister | 10 of 10 | NMA | FCR | 5.5 | PD | 450 | 1 | yes, stage 3 skin | moderate | progressive disease |
| 4 | MUD | 10 of 10 | NMA | FCRA | 10.4 | CRi w/ MRD | 271 | 3 | no | severe | GVHD and infection |
| 5 | sister | 9 of 10 (1 mismatch at DQB1) | NMA | FCR | 6.2 | PR | 165 | 5 | no | no | progressive disease and infection |
| 6 | sister | 10 of 10 | NMA | FCR | 2.4 | SD | 1072 | 0 | yes, stage 3 skin | no | progressive disease, Richter's syndrome |
| 7 | sister | 10 of 10 | NMA | FCR | 5.5 | PR | 396 | 1 | yes, stage 3 GI | no | progressive disease and infection |
| 8 | sister | 10 of 10 | NMA | FCR | 3.6 | PR | 177 | 2 | no | no | progressive disease and infection |
| 9 | brother | identical by serology and PCR-SSP | RIC | FA + M^#^ | 6.3 | CRi w/ MRD | 406 | 1 | no | moderate | FTT and infection |
| 10 | brother | 10 of 10 | NMA | FCRI | 1.1 | PR | 619 | 3 | no | no | alive |
| 11 | UCB | 10 of 10 | NMA | FCRT | 7.4 | PR | 690 | 0 | no | severe | progressive disease and infection |
| 12 | MUD | 10 of 10 | NMA | FCRA | 7.9 | CR | 2320 | 0 | no | severe | unknown |
| 13 | MUD | 10 of 10 | RIC | FR + M | 6.0 | SD | 3124 | 0 | no | moderate | progressive disease |
| 14 | MUD | 10 of 10 | RIC | FR + M | 11.3 | PR | 3955 | 0 | yes, stage 3 skin | moderate | unknown |
| 15 | sister | 10 of 10 | NMA | FCR | 18.1 | PR | 4687 | 0 | no | no | alive |
| 16 | sister | 10 of 10 | NMA | FCR | 6.4 | CR | 4500 | 0 | yes, stage 3 skin | no | lost to follow up |
| 17 | MUD | 9 of 10 (1 mismatch at C locus) | NMA | FCR | 3.1 | PR | 4808 | 0 | yes, stage 3 skin | moderate | lost to follow up |
| 18 | MUD | 10 of 10 | MA | BEAM+AR | 8.1 | PR | 3311 | 0 | no | no | lost to follow up |
| 19 | MUD | 10 of 10 | NMA | FCRI | 13.9 | PR | 2521 | 0 | no | no | lost to follow up |
| 20 | brother | 10 of 10 | NMA | FCR | 1.6 | CR | 3821 | 0 | no | no | lost to follow up |
| 21 | MUD | 10 of 10 | NMA | FCR | 5.9 | PR | 2662 | 0 | no | moderate | lost to follow up |
| 22 | sister | 10 of 10 | RIC | FR + M | 7.8 | SD | 426 | 0 | no | severe | steroid refractory GVHD and infection |
| 23 | sister | 10 of 10 | RIC | F + M | 9.4 | PR | 463 | 0 | yes, stage 3 skin | severe | steroid refractory GVHD and infection |
| 24 | brother | 10 of 10 | NMA | R + I | 7.5 | PR | 3591 | 0 | yes, stage 3 skin | no | unknown |

### Slide 3
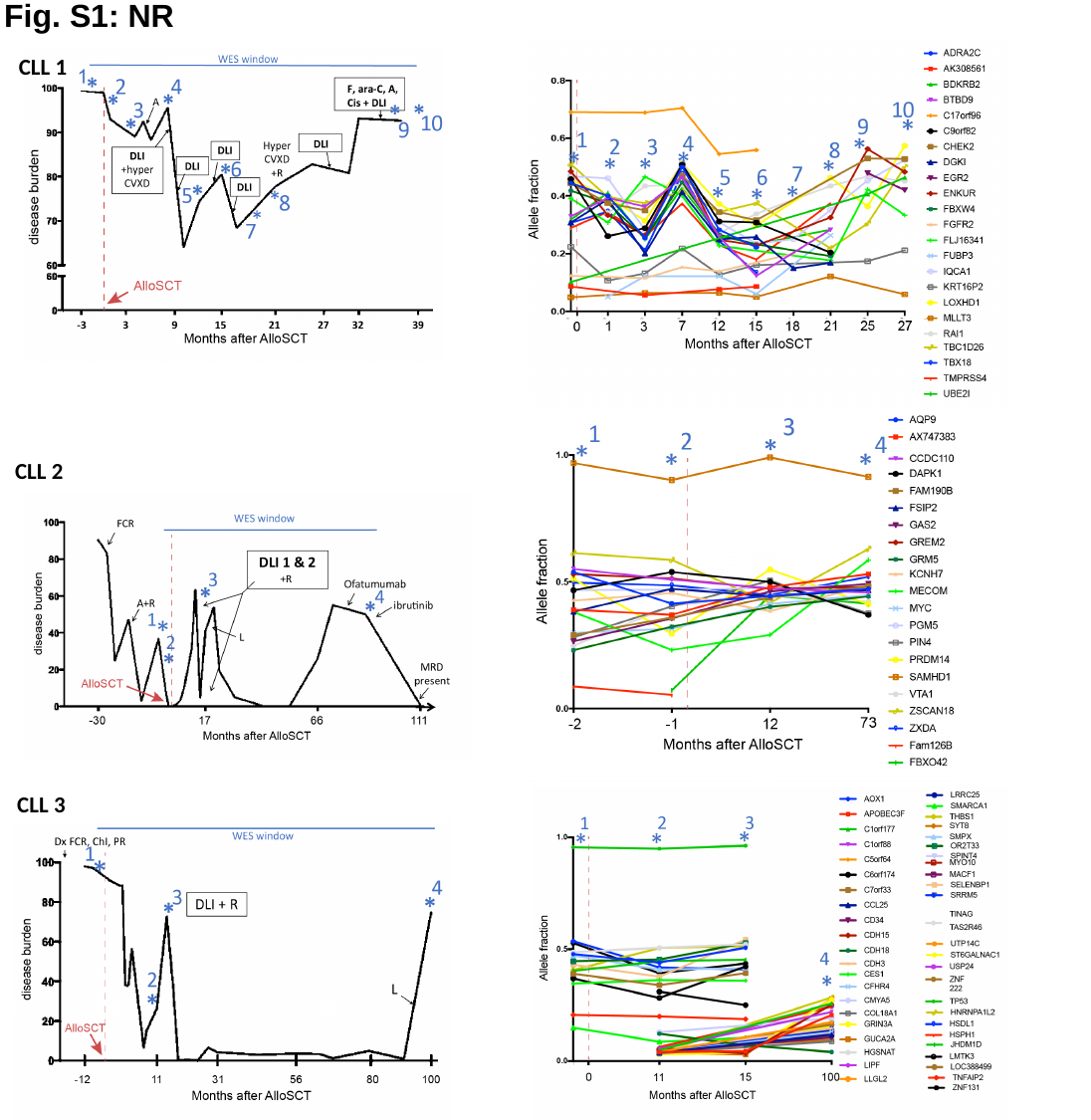

Fig. S1: NR
CLL 1
CLL 2
CLL 3

### Slide 4
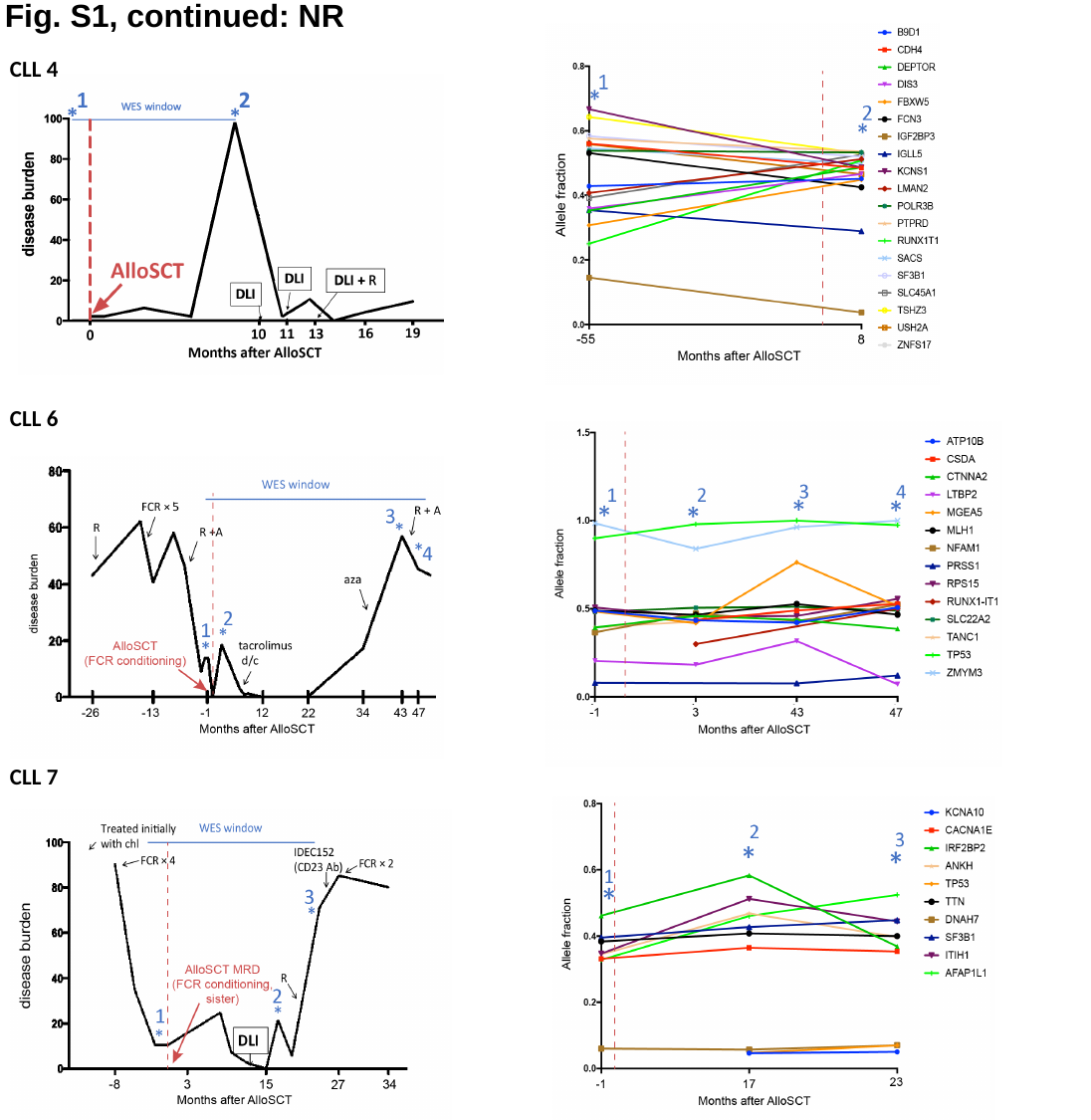

Fig. S1, continued: NR
CLL 4
CLL 6
CLL 7

### Slide 5
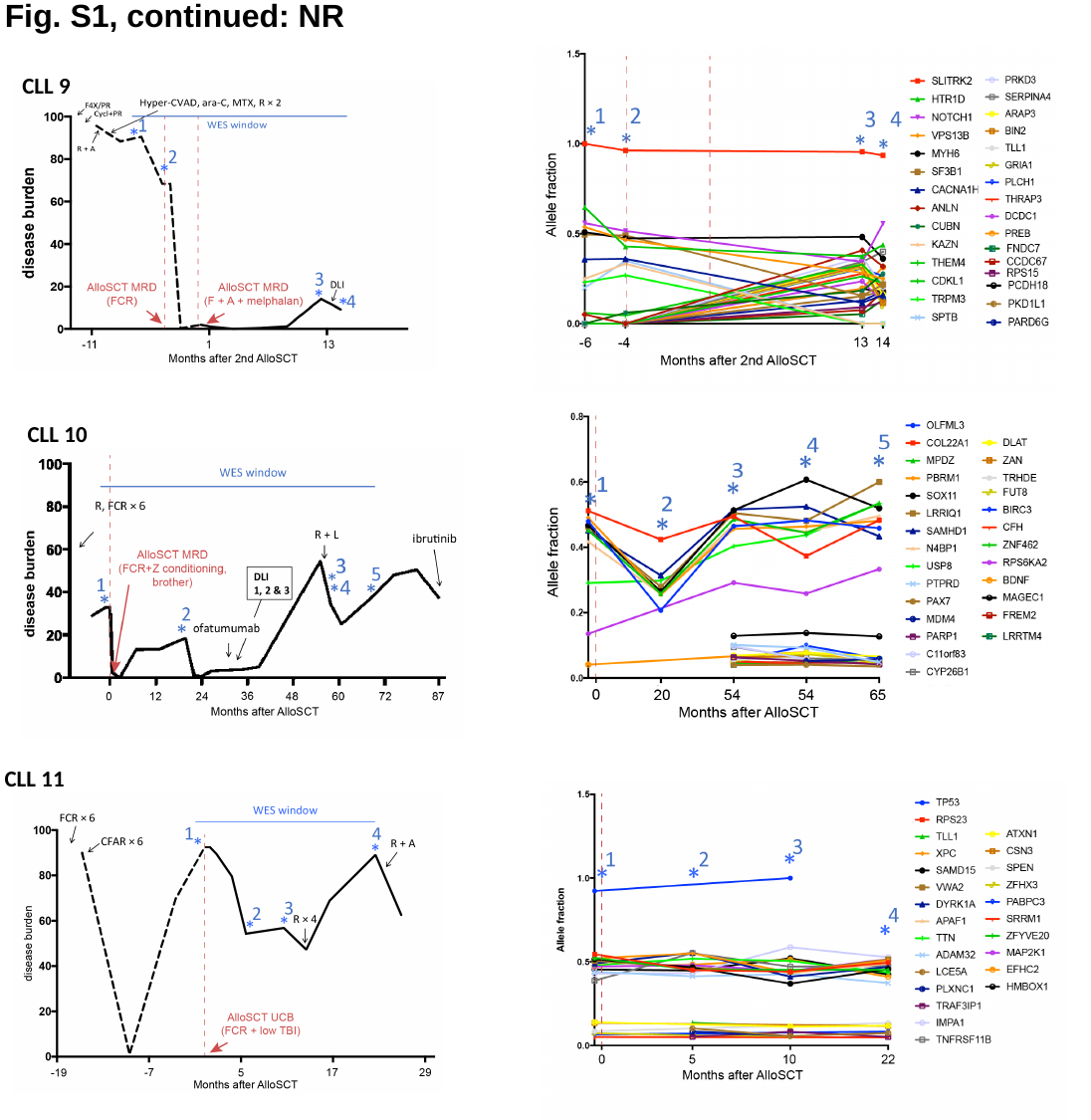

Fig. S1, continued: NR
CLL 9
CLL 10
CLL 11

### Slide 6
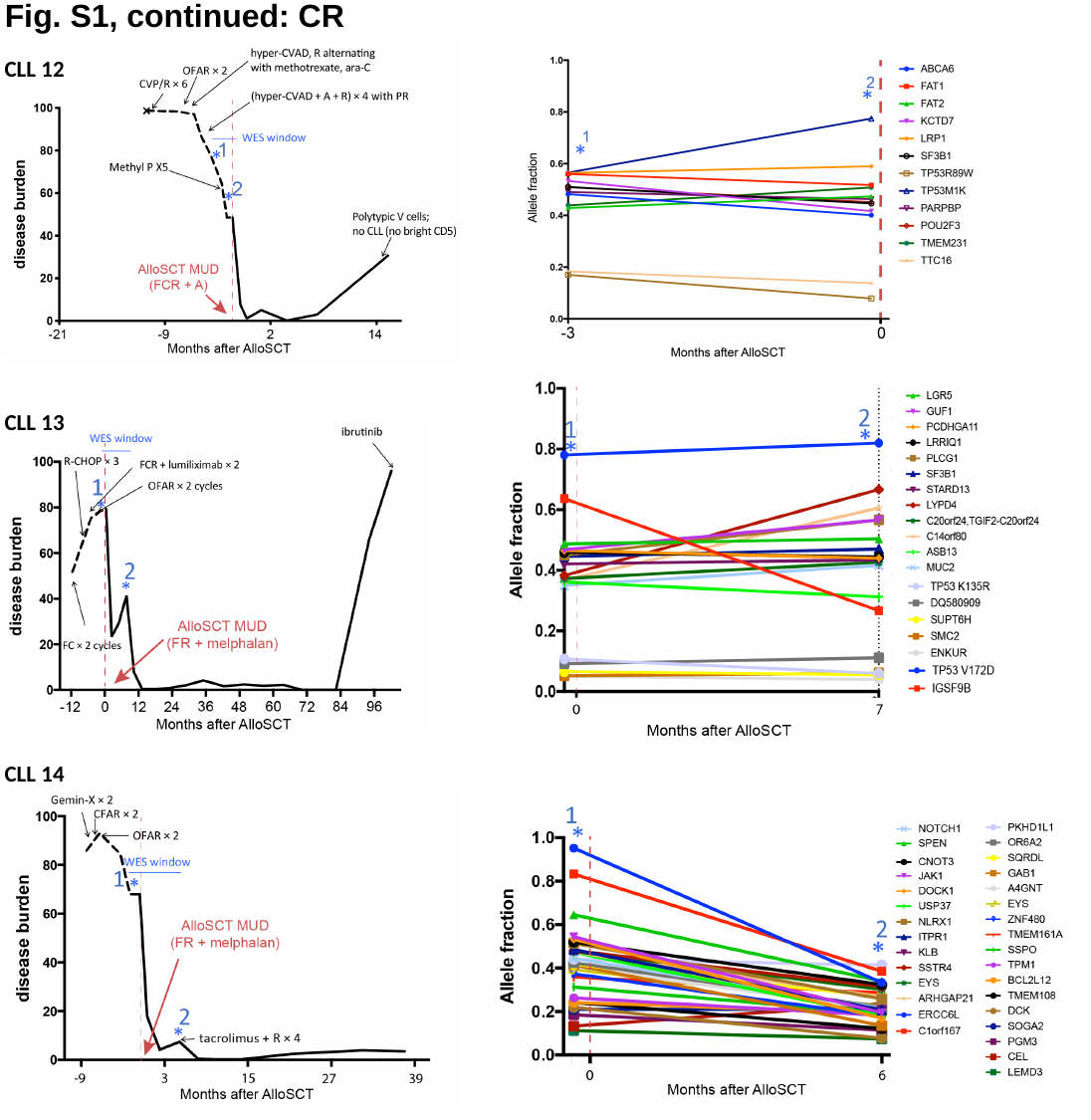

Fig. S1, continued: CR
CLL 12
CLL 13
CLL 14

### Slide 7
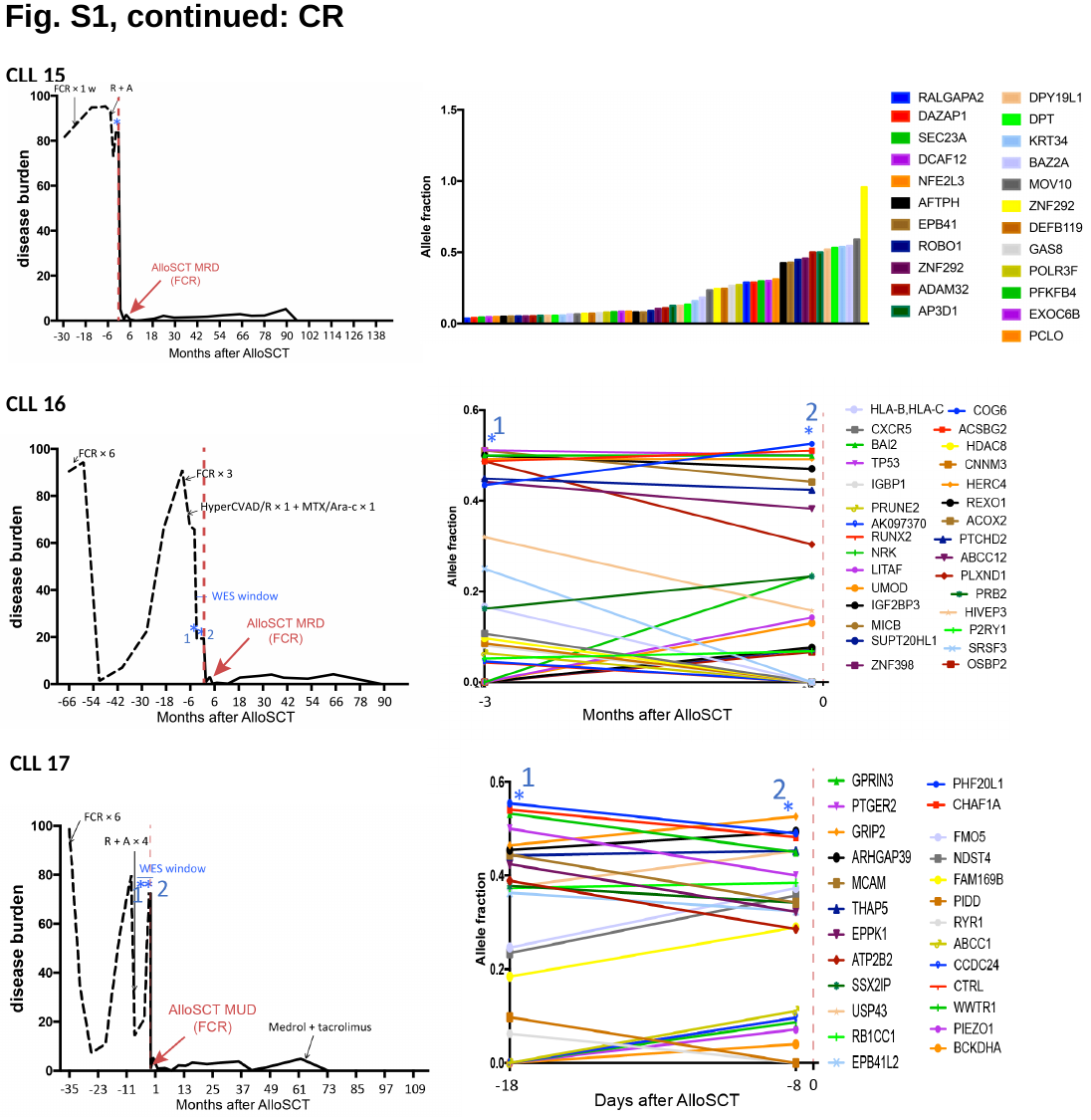

Fig. S1, continued: CR
CLL 15
CLL 16
CLL 17

### Slide 8
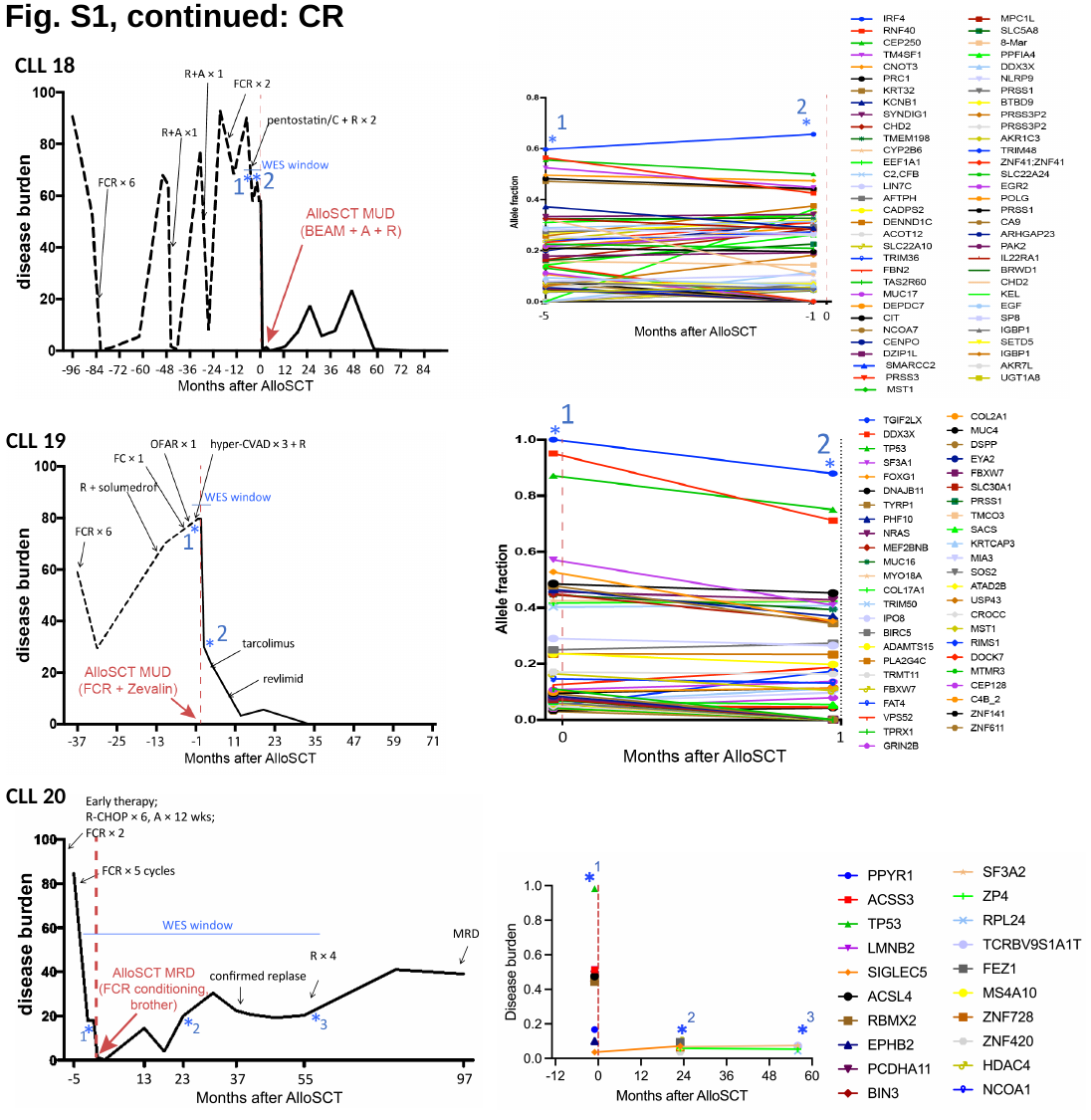

Fig. S1, continued: CR
CLL 18
CLL 19
CLL 20

### Slide 9
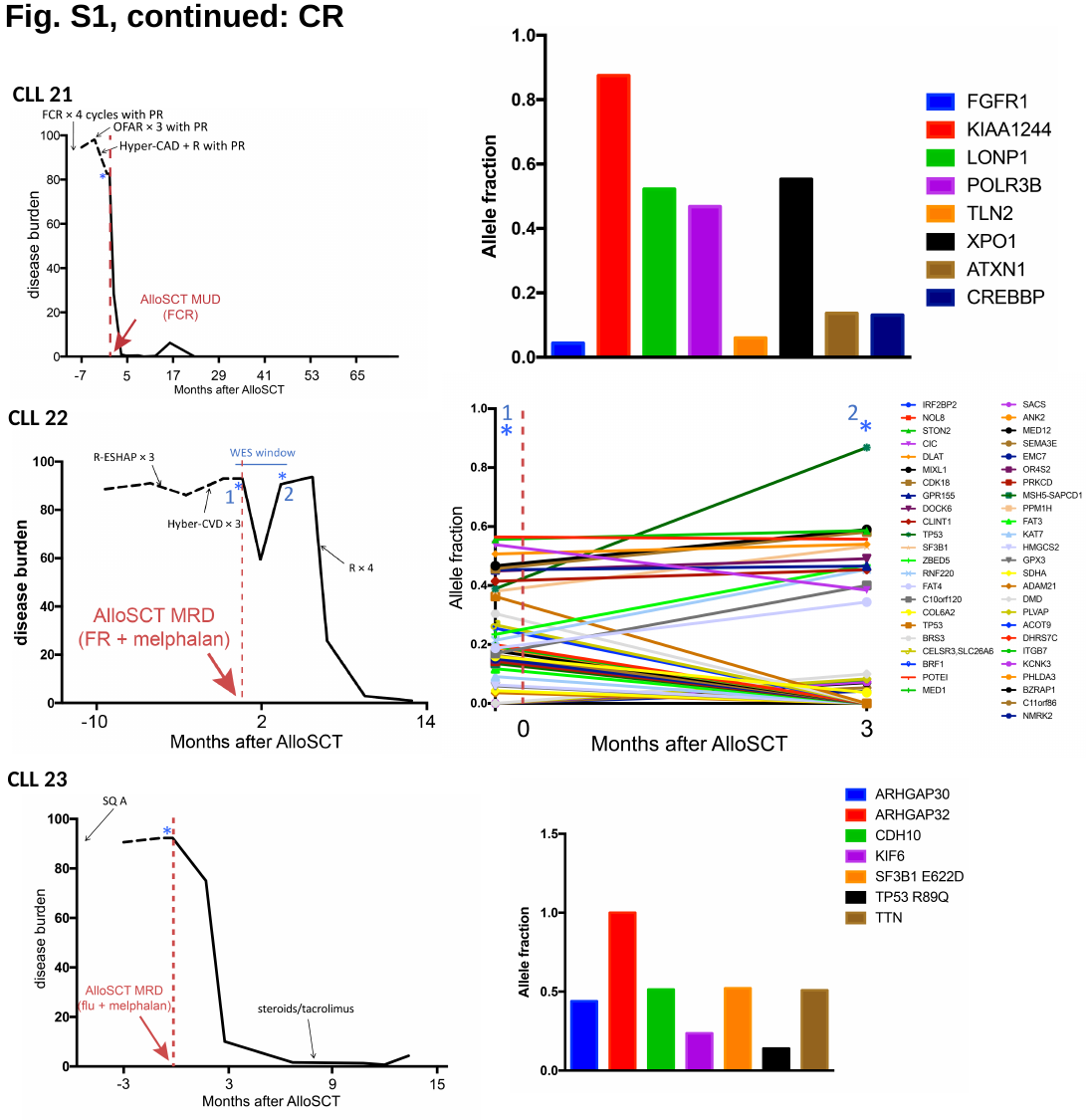

Fig. S1, continued: CR
CLL 21
CLL 22
CLL 23

### Slide 10
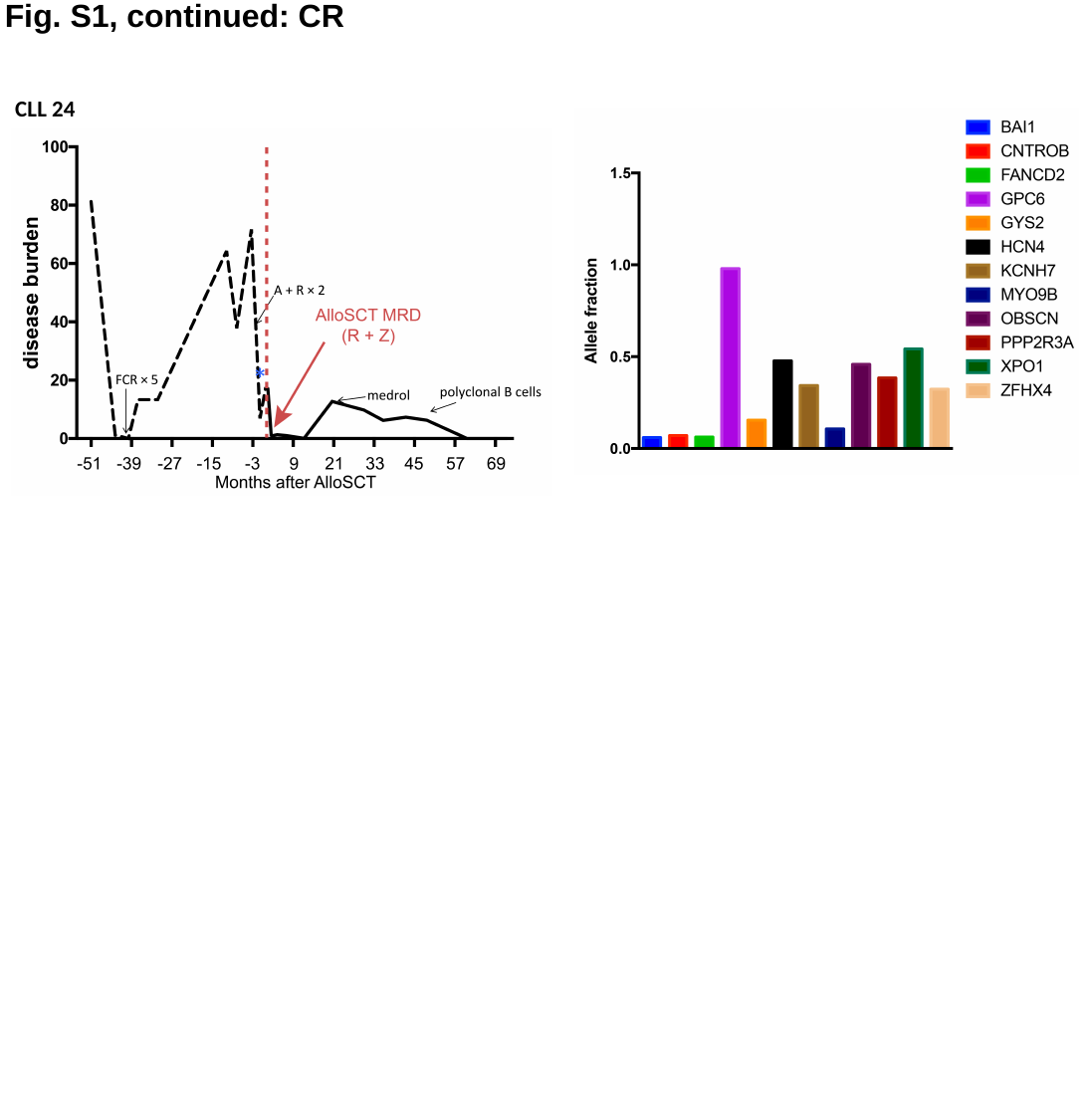

Fig. S1, continued: CR
CLL 24

### Slide 11
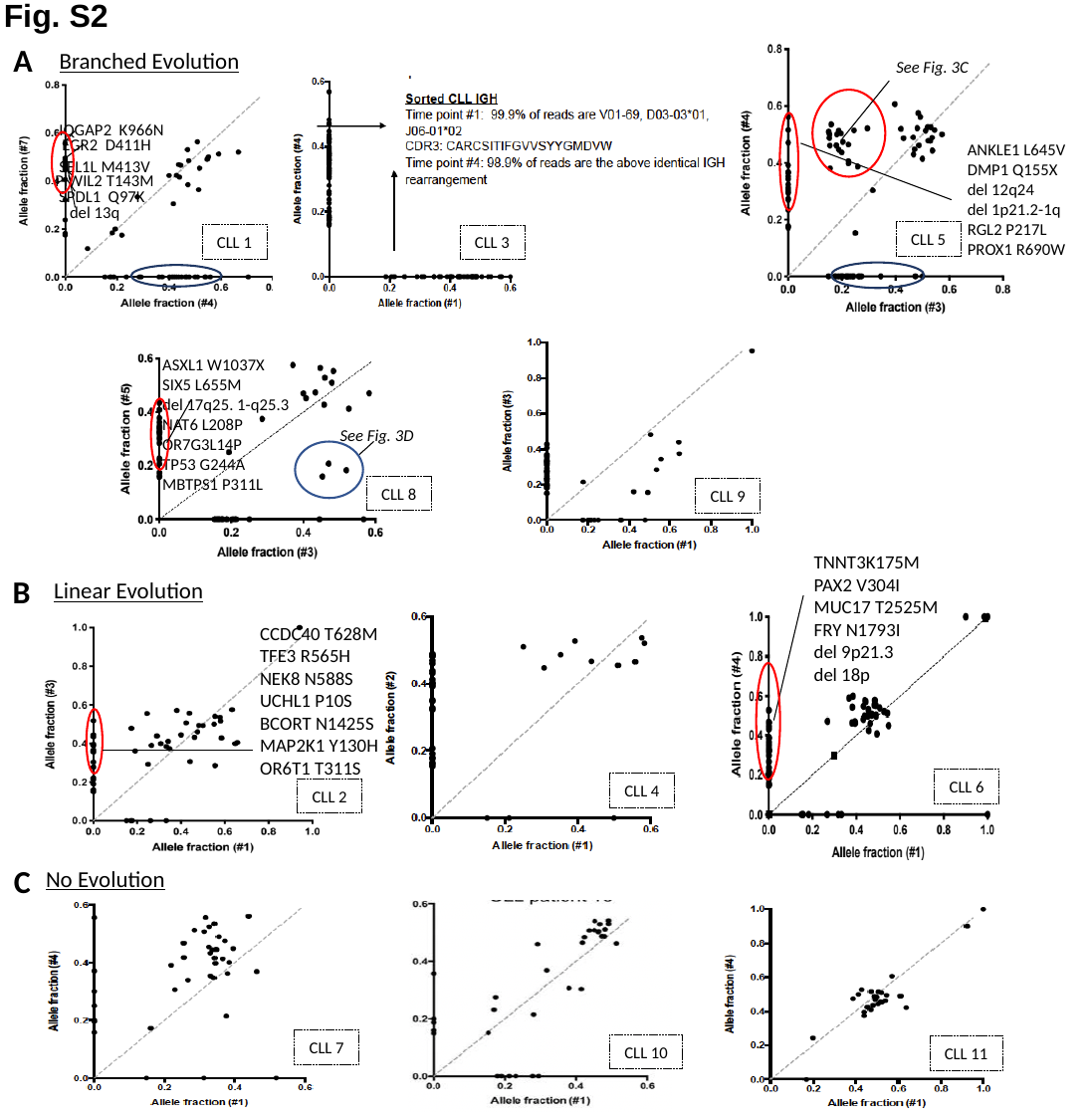

Fig. S2
A
Branched Evolution
IQGAP2 K966N
EGR2 D411H
SEL1L M413V
PIWIL2 T143M
SPDL1 Q97K
del 13q
CLL 1
ANKLE1 L645V
DMP1 Q155X
del 12q24
del 1p21.2-1q
RGL2 P217L
PROX1 R690W
CLL 5
See Fig. 3C
CLL 3
See Fig. 3D
ASXL1 W1037X
SIX5 L655M
del 17q25. 1-q25.3
NAT6 L208P
OR7G3L14P
TP53 G244A
MBTPS1 P311L
CLL 8
CLL 9
TNNT3K175M
PAX2 V304I
MUC17 T2525M
FRY N1793I
del 9p21.3
del 18p
B
Linear Evolution
CCDC40 T628M
TFE3 R565H
NEK8 N588S
UCHL1 P10S
BCORT N1425S
MAP2K1 Y130H
OR6T1 T311S
CLL 6
CLL 4
CLL 2
C
No Evolution
CLL 7
CLL 10
CLL 11

### Slide 12
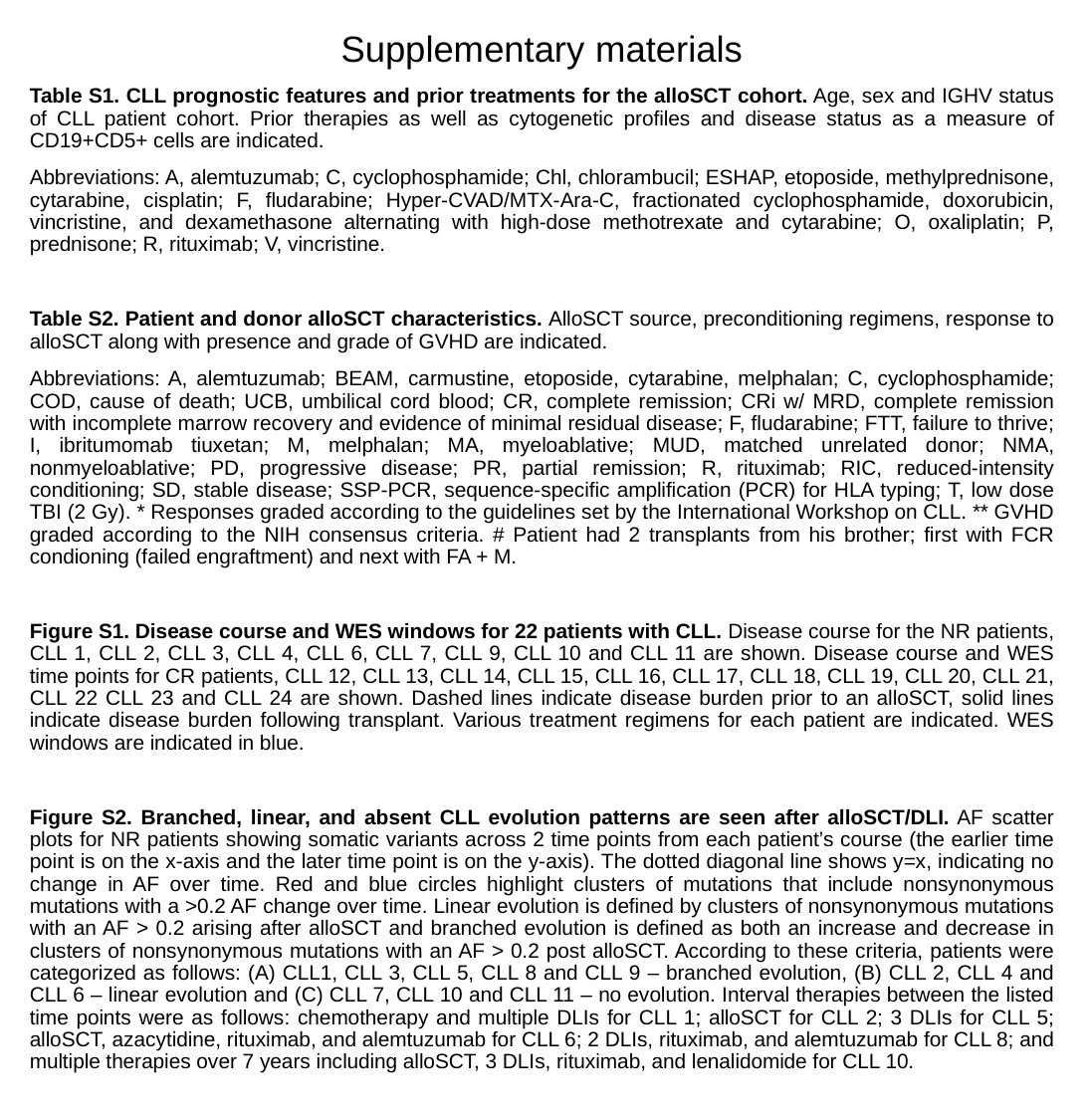

Supplementary materials
Table S1. CLL prognostic features and prior treatments for the alloSCT cohort. Age, sex and IGHV status of CLL patient cohort. Prior therapies as well as cytogenetic profiles and disease status as a measure of CD19+CD5+ cells are indicated.
Abbreviations: A, alemtuzumab; C, cyclophosphamide; Chl, chlorambucil; ESHAP, etoposide, methylprednisone, cytarabine, cisplatin; F, fludarabine; Hyper-CVAD/MTX-Ara-C, fractionated cyclophosphamide, doxorubicin, vincristine, and dexamethasone alternating with high-dose methotrexate and cytarabine; O, oxaliplatin; P, prednisone; R, rituximab; V, vincristine.
Table S2. Patient and donor alloSCT characteristics. AlloSCT source, preconditioning regimens, response to alloSCT along with presence and grade of GVHD are indicated.
Abbreviations: A, alemtuzumab; BEAM, carmustine, etoposide, cytarabine, melphalan; C, cyclophosphamide; COD, cause of death; UCB, umbilical cord blood; CR, complete remission; CRi w/ MRD, complete remission with incomplete marrow recovery and evidence of minimal residual disease; F, fludarabine; FTT, failure to thrive; I, ibritumomab tiuxetan; M, melphalan; MA, myeloablative; MUD, matched unrelated donor; NMA, nonmyeloablative; PD, progressive disease; PR, partial remission; R, rituximab; RIC, reduced-intensity conditioning; SD, stable disease; SSP-PCR, sequence-specific amplification (PCR) for HLA typing; T, low dose TBI (2 Gy). * Responses graded according to the guidelines set by the International Workshop on CLL. ** GVHD graded according to the NIH consensus criteria. # Patient had 2 transplants from his brother; first with FCR condioning (failed engraftment) and next with FA + M.
Figure S1. Disease course and WES windows for 22 patients with CLL. Disease course for the NR patients, CLL 1, CLL 2, CLL 3, CLL 4, CLL 6, CLL 7, CLL 9, CLL 10 and CLL 11 are shown. Disease course and WES time points for CR patients, CLL 12, CLL 13, CLL 14, CLL 15, CLL 16, CLL 17, CLL 18, CLL 19, CLL 20, CLL 21, CLL 22 CLL 23 and CLL 24 are shown. Dashed lines indicate disease burden prior to an alloSCT, solid lines indicate disease burden following transplant. Various treatment regimens for each patient are indicated. WES windows are indicated in blue.
Figure S2. Branched, linear, and absent CLL evolution patterns are seen after alloSCT/DLI. AF scatter plots for NR patients showing somatic variants across 2 time points from each patient’s course (the earlier time point is on the x-axis and the later time point is on the y-axis). The dotted diagonal line shows y=x, indicating no change in AF over time. Red and blue circles highlight clusters of mutations that include nonsynonymous mutations with a >0.2 AF change over time. Linear evolution is defined by clusters of nonsynonymous mutations with an AF > 0.2 arising after alloSCT and branched evolution is defined as both an increase and decrease in clusters of nonsynonymous mutations with an AF > 0.2 post alloSCT. According to these criteria, patients were categorized as follows: (A) CLL1, CLL 3, CLL 5, CLL 8 and CLL 9 – branched evolution, (B) CLL 2, CLL 4 and CLL 6 – linear evolution and (C) CLL 7, CLL 10 and CLL 11 – no evolution. Interval therapies between the listed time points were as follows: chemotherapy and multiple DLIs for CLL 1; alloSCT for CLL 2; 3 DLIs for CLL 5; alloSCT, azacytidine, rituximab, and alemtuzumab for CLL 6; 2 DLIs, rituximab, and alemtuzumab for CLL 8; and multiple therapies over 7 years including alloSCT, 3 DLIs, rituximab, and lenalidomide for CLL 10.
